# A potential role of altered SIgA-targeting of gut microbiota in long-term dysbiosis in pediatric solid organ transplant recipients

**DOI:** 10.1101/2025.08.11.668420

**Authors:** Kora Schulze, Imeke Goldschmidt, Anette Melk, Martin Böhne, Sabrina Woltemate, Matthias Ballmaier, Svea Kleiner, Elena Lehmann, Miriam Kramer, Marius Vital

**Affiliations:** Institute for Medical Microbiology and Hospital Epidemiology, Hannover Medical School, Carl-Neuberg-Strasse 1, 30625, Hannover, Germany; Department of Pediatric Liver, Kidney and Metabolic Diseases, Division of Pediatric Hepatology and Liver Transplantation, Hannover Medical School, Carl-Neuberg-Strasse 1, 30625, Hannover, Germany; Department of Pediatric Liver, Kidney and Metabolic Diseases, Hannover Medical School, Carl-Neuberg-Strasse 1, 30625, Hannover, Germany; Department of Pediatric Cardiology and Intensive Care, Hannover Medical School, Carl-Neuberg-Strasse 1, 30625, Hannover, Germany; Central Research Facility Cell Sorting, Hannover Medical School, Carl-Neuberg-Strasse 1, 30625, Hannover, Germany

**Keywords:** microbiota, IgA, solid organ transplantation, immunosuppression, metagenomics

## Abstract

**Background:** Composition of gut microbiota (GM) is altered in solid organ transplantation (SOT) patients, where the degree of dysbiosis is associated with long-term survival. Factors governing dysbiosis are poorly understood with immunosuppression therapy believed to be involved in altering GM composition in those patients, either directly or via the immune system. At the interface stands secretory (S)IgA, however, little is known on its role governing dysbiosis in the context of SOT. In this study, we performed quantitative metagenomic analyses of GM accompanied by SIgA sequencing in 48 pediatric SOT recipients (age = 10.6 ± 4.7 years) receiving either a heart (HTR, n=10), a kidney (KTR, n=11) or a liver (LTR, n=27) and compared results to healthy controls (HC, n=16).

**Results:** We confirmed compositional and functional dysbiosis in SOT patients that broadly clustered into two groups, where cluster 1 (n=23) comprised most LTR patients and was closer to HC compared with cluster 2 (n=24) that exhibited stronger dysbiosis and included most HTR and KTR patients. The degree of dysbiosis was associated with Tacrolimus (TAC) levels. Overall, patients exhibited higher SIgA levels than HC, along with an increased percentage of bacteria targeted and altered target spectra. Furthermore, altered SIgA responses were associated with the degree of dysbiosis and were especially increased in cluster 2, in particular in HTR patients. A mechanistic model connecting immunosuppression, GM composition and SIgA targeting is proposed.

**Conclusion:** Our study suggests that altered SIgA responses play an important role for GM alterations observed in SOT patients. It opens new therapeutic angles to combat GM dysbiosis and associated long-erm complications.

## Introduction

Solid organ transplantation (SOT) is a final therapy option in patients suffering from severe diseases with irreversible and life-endangering progression that cannot be treated otherwise.^1^ Subsequent immunosuppression is required throughout life to prevent graft rejection. The dose and regimen of immunosuppressants is adapted based on the organ transplanted, the time since transplantation, the age of the patients as well as individual variability which necessities the requirement of a tight therapeutic monitoring also after surgery. While strong and often repeated antibiotic treatments before and short-term after SOT are known to cause gut microbiota (GM) dysbiosis,^2^ there is increasing evidence that immunosuppressants overall decrease the bacterial richness and diversity, potentially causing an altered GM composition also several years post-transplantation.^2–4^ Moreover, studies showed that GM are capable of modulating drug pharmacokinetics, for example by altering transporter expression or metabolizing the drug itself, highlighting the bidirectional relationship between immunosuppression and GM.^5,6^ Evidence regarding effects of SOT and immunosuppression on GM is scarce and very heterogeneous regarding the study design and the characteristics of the analyzed patient cohort. Furthermore, the extrapolation of results from adult to children is limited, as pediatric SOT recipients usually have different underlying diseases as indication for transplantation, often require a more dynamic immunosuppressant regimen and show a higher risk and frequency of infection and transplant-related complications.^7–10^ As observed by us and others, GM dysbiosis in pediatric SOT recipients is associated with short- and long-term health outcomes, including an increased risk for infection, graft health and all-cause mortality, highlighting the importance to combat GM dysbiosis in those patients.^2,11,12^

The interaction between the host immune system and GM is regulated to a large extend via the secretory immunoglobulin A (SIgA). SIgA is the most abundant immunoglobulin isotype in mammals and predominantly located at mucosal surfaces, where its main function is the exclusion of pathogens.^16,17^ However, recent years revealed that it is also targeting commensal bacteria, thereby regulating their colonization, growth and motility.^18,19^ The underlying mechanisms by which SIgA exerts its various effects on GM are hypothesized to be the result of SIgA affinity maturation and consequent differences in binding to the respective bacteria, as outlined in several reviews on this topic.^20,21^ SIgA-targeting of GM and its impact on host immunity is also of great importance for child development. During the first days of life, maternal breastmilk is the only source of SIgA for the neonate, eliciting an important role in the early shaping of the infants GM.^22,23^ Ding et al.^24^ reviewed SIgA-targeting patterns and consequent changes in microbiota composition during child development, as well as factors influencing this relation, highlighting the role of SIgA in the maturation of the infants’ immune system and connecting altered SIgA-targeting to diseases later in life, such as allergies, diarrhea and colitis. The association between altered SIgA-targeting and disease was also intensively investigated in adults, where it has often been associated with infection and inflammation along with overall GM dysbiosis.^19,25^

Immunosuppression in SOT recipients targets immune cells that are also involved in the synthesis and maturation of IgA, namely B and T cells, however, whether those patients show an altered IgA expression as well as a subsequent altered SIgA-targeting of GM that eventually contributes to the observed dysbiosis, is unknown. To test this hypothesis, we investigated the SIgA-targeting of GM in pediatric SOT recipients under stable immunosuppression and compared the results to an age-matched healthy cohort providing the first study to investigate the effect of immunosuppression on SIgA targeting of GM and its relation to dysbiosis in patients after SOT. We compared different organ transplants (heart, liver, kidney) as well as different immunosuppressant regimens from a major German transplant center minimizing regional variability of GM and differences in surgery and post-surgery treatment.

## Results

### Description of the study cohort

Stool samples from 48 patients (age = 10.4 ± 4.74 years) and 16 age-matched healthy controls (HC; age = 10.2 ± 4.81 years) were analyzed. All patients had undergone SOT, with n=10 receiving a kidney (KTR), n=11 receiving a heart (HTR) and n=27 receiving a liver transplant (LTR). Samples were collected on average 6.80 ± 4.19 years after transplantation. A detailed overview of patient characteristics and medication is provided in **Table 1**. The majority of patients (n=34) got Tacrolimus (TAC) as the primary immunosuppressant, whereas the other 14 patients received Cyclosporine A (CSA). LTR were treated with either TAC or CSA as monotherapy, whereas KTR or HTR were also administered steroids; most of latter patients (n=15) additionally got a purine-analogue or mTOR inhibitor as third immunosuppressant.

**Table 1.** Characteristics of patients.

|  |  | Patients (n=48) | HC (n=16) |
| --- | --- | --- | --- |
| Age (years) |  | 10.6 ± 4.7 | 10.2 ± 4.8 |
| Organ transplanted (n) |  |  |  |
|  | liver | 27 |  |
|  | heart | 11 |  |
|  | kidney | 10 |  |
| Time since transplant (years) |  | 6.8 ± 4.2 |  |
| Immunosuppressant |  |  |  |
| a. total number |  |  |  |
|  | n=1 | 27 |  |
|  | n=2 | 6 |  |
|  | n=3 | 15 |  |
| b. type |  |  |  |
|  | Tacrolimus (TAC) | 34 |  |
|  | Cyclosporine A (CSA) | 14 |  |
|  | Everolimus | 9 |  |
|  | Sirolimus | 1 |  |
|  | Mycophenolat (MMF) | 4 |  |
|  | Azathioprin | 1 |  |
|  | Prednisolon | 21 |  |
| c. level | Tacrolimus [μl/l] | 5.1 ± 2.8 |  |
|  | CSA [μg/l] | 44.1 ± 15.6 |  |
|  | Everolimus [μl/l] | 3.5 ± 1.1 |  |

### Patients clustered into two groups based on GM composition

Metagenomic analysis based on species level grouped samples into two distinct clusters (C1 and C2) as determined by k-means clustering analysis based on Bray-Curtis dissimilarities (BC) (**Figure 1A**, **Supplementary figure S1**). C1 included all healthy samples and the majority of samples from LTR; for all following analysis, C1 is only referring to patients within that cluster (n=23), whereas samples derived from healthy children were treated as a separate (control) group (HC, n=16). Samples in C2 (n=24) comprised most KTR and HTR. To compare GM dysbiosis between patient groups, the BC distance of each patient sample was calculated to the average BC dissimilarity in HC (BC to HC). C2 displayed a significantly higher dysbiosis expressed as BC to HC, namely 0.835 ± 0.067, compared with samples from C1 (0.711 ± 0.048) (**Figure 1A, D**). There was no difference in the age distribution between any cluster and HC (**Figure 1C**), however, patients in C2 trended towards a lower time since transplant (ΔTx, 5.82 ± 3.91 years) compared with C1 (8.08 ± 4.14 years, *p=0.0918*). Consequently, time since transplant was included as a fixed effect variable in all further analyses. There was a trend towards higher TAC levels in C2 (5.77 ± 2.62 µl/l) compared with C1 (4.42 ± 2.92 µl/l, *p=0.0819*), but this observation disappeared when organ was taken into account. When comparing organs, TAC levels were significantly higher in both KTR (6.32 ± 4.33 µl/l, *p=0.0414*) and HTR (7.43 ± 2.63 µl/l, *p=0.0005*) compared with LTR (3.77 ± 1.32 µl/l). No difference in average CSA levels between clusters or organs was observed. One LTR sample (“L07”) was located far outside the two identified clusters displaying a strong dominance of the genus Klebsiella (77.7 %) (**Figure 1B**) and this sample was, hence, excluded from all further statistical analyses.

**Figure 1.**
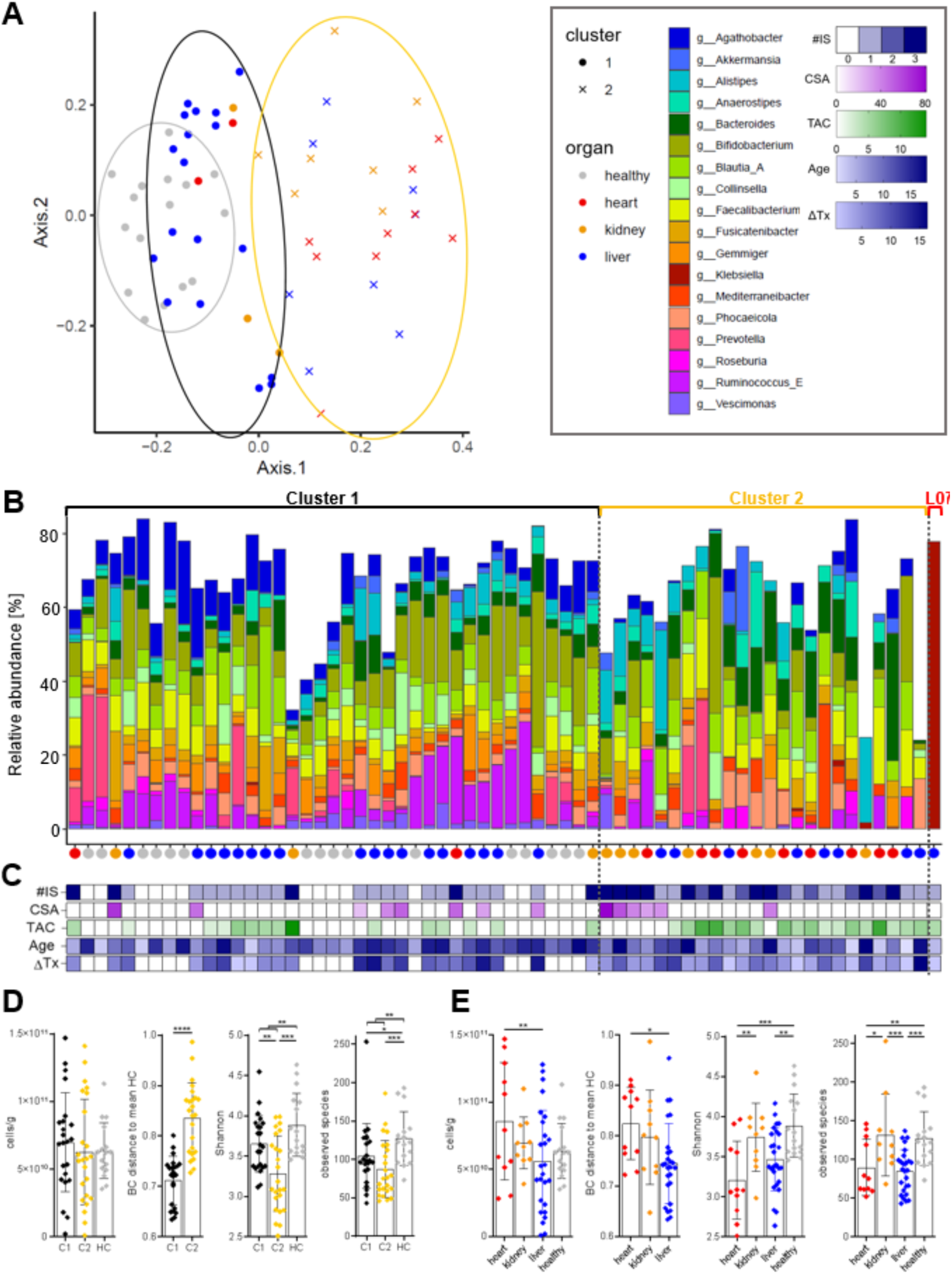
Comparison of bacterial communities between patients and healthy controls (HC). Panel **A** depicts the metric multidimensional scaling analysis of all samples based on Bray Curtis dissimilarities (BC) of metagenomics sequencing data on species level. Clustering was done by the k-means algorithm with colored circles indicating respective clusters (black = C1, gold = C2, grey = HC). Samples are additionally colored according to the respective organ group. Panel **B** shows the relative abundance of major genera (mean relative abundance >1%) for each sample grouped by the identified clusters shown in **A**. Colored dots below the x-axis indicate the respective organ group of each sample. The sample L07 was allocated outside the identified two clusters and is shown separately. In panel **C** the information on administered immunosuppressants, blood levels of Tacrolimus (TAC) and Cyclosporine A (CSA), age and time since transplantation (ΔTx) for each sample is given; white color indicates no data. The distribution of bacterial cell counts, the BC distance in each sample to the average BC dissimilarity in HC (representing the degree of dysbiosis) (BC to HC), the Shannon index and number of observed species per cluster and organ group are given in panel **D** and **E**, respectively. * - p<0.05, ** - p<0.01, *** - p<0.001, **** - p<0.0001.

The total bacterial cell count was similar between the two clusters but was significantly higher in HTR (8.57 x 10^10^ ± 4.37 x 10^10^ cells/g) compared with LTR (5.62 x 10^10^ ± 3.79 x 10^10^ cells/g) (**Figure 1D, E**). Bacterial cell counts were not different in any organ group when compared with HC. HTR had a larger BC to HC (0.824 ± 0.072) compared with LTR (0.745 ± 0.079) (**Figure 1E**), whereas KTR did not significantly differ in BC to HC (0.797 ± 0.094) from the two other organ groups. C2 was characterized by a significantly reduced Shannon index (3.28 ± 0.46) compared with HC (3.89 ± 0.39) and C1 (3.65 ± 0.35) (**Figure 1D**). A similar pattern was observed when comparing the number of observed species between clusters (**Figure 1D**). On the organ level, HTR had a significantly reduced diversity compared with HC as indicated by a lower Shannon index (HTR: 3.21 ± 0.49; HC: 3.89 ± 0.39) as well as a lower number of observed species (HTR: 89.9 ± 52.9; HC: 127 ± 34.8). Similar results were found in LTR, whereas KTR did not show differences in diversity compared with HC (**Figure 1E**).

GM composition on the phylum level was characterized by an increased abundance in Verrucomicrobiota in patients compared with HC, however only as a trend (**Figure 2A**). Analyses of major genera (mean relative abundance >1 %) showed a reduced relative abundance of the taxa Agathobacter in patients compared with HC (**Figure 2B**). On species level, *Agathobacter rectalis* was significantly lower abundant in patients. Additionally there was a trend towards a lower abundance of and a trend towards a lower relative abundance of Gemmiger and Ruminococcus_E in patients compared with HC. When comparing clusters, C2 was characterized by a significantly lower relative abundance of bacteria belonging to the phylum Actinomycetota and a higher abundance of bacteria belonging to the phylum Bacillota_A compared with HC. Furthermore, C2 was characterized by reduced abundances of the genera Agathobacter, Gemmiger and Ruminococcus_E compared with HC. The dominant species of those genera (relative abundance of >0.1 %) were *Agathobacter rectalis, Gemmiger qucibialis* and *Ruminococcus_E bromii*, respectively, that were also lower abundant in C2 compared with HC, albeit the latter only as a trend. Furthermore, C2 trended towards lower relative abundances of Bifidobacterium and Vescimonas, and a higher relative abundance of Bacteroides (**Figure 2B**). Regarding the former, several highly abundant species, namely, *B. adolescentis, B. catenulatum* and *B. infantis*, were reduced in C2 compared with HC (data not shown). None of the major genera had a significantly different relative abundance in C1 compared with HC (**Figure 2B**).

**Figure 2.**
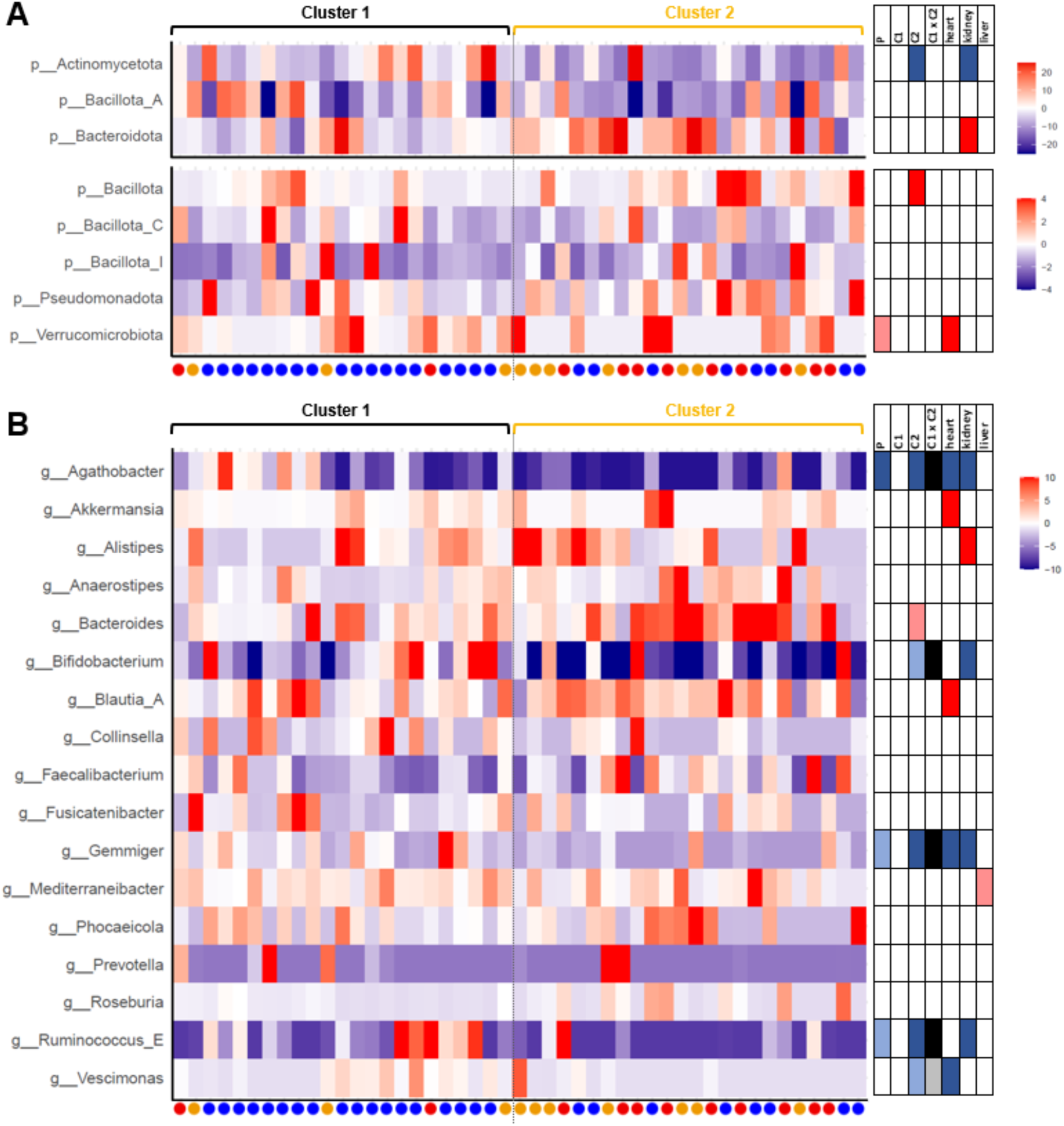
Differences in the relative abundance of taxa in patient samples compared with healthy controls (HC, mean relative abundance). Taxa with a mean relative abundance >1% are shown on the phylum level (panel **A**) and the genus level (panel **B**). Samples are ordered according to Figure 1B. Dots below the x-axis indicate the respective organ group of each sample. The tables on the right give information about significant differences for the respective group with red and blue colors indicating a higher and lower relative abundance, respectively, compared with HC. Dark colors indicate significant differences (*lfdr*<0.05), whereas light colors indicate trends (*lfdr*<0.1). For comparison between clusters (C1xC2), black and grey color indicate significant and trending lower abundance, respectively, of a taxon in C2 compared with C1.

Analyses on the organ level showed higher abundances of the phylum Verrucomicrobiota and the genera Akkermansia and Blautia_A and lower abundances of Agathobacter, Gemmiger and Vescimonas in HTR compared with HC (**Figure 2B**). Bacteria belonging to the genus Alistipes were more abundant in KTR compared with HC whereas Agathobacter, Bifidobacterium, Gemmiger and Ruminococcus_E were lower abundant in this group. Mediterraneibacter was more abundant in LTR, albeit only as a trend. Overall, HTR had the largest number of differently abundant taxa of all organ groups compared with HC. GM of LTR resembled most closely the composition found in healthy controls with lowest number of taxa being differently abundant, which is in accordance with overall clustering analyses shown above.

Functional analyses of gut GM confirmed the differences observed between clusters based on taxonomic composition. C2 had a significantly higher BC to HC (0.182 ± 0.031) compared with C1 (0.146 ± 0.025) when based on abundances of KEGG orthologues (**Supplementary Figure S2A**). A significant higher BC to HC in C2 compared to C1 was also observed based on the abundances of CAZymes (0.234 ± 0.056 vs. 0.171 ± 0.032) (**Supplementary Figure S2B**). Overall, the functional potential in samples derived from C2 was characterized by many abundance differences compared with HC, where 46.9 % of all KEGG orthologues, 58.8 % of all KEGG modules and 38.0 % of all CAZymes were higher abundant in C2, whereas 4.7 %, 0.7 % and 9.1 %, respectively, were more abundant in HC (**Supplementary Figure S2C**). In contrast, there were no significant differences in the relative abundance of any functions between C1 and HC, further supporting the similarity between samples of this cluster with HC also on the functional level.

### TAC had a larger impact on GM composition than CSA

Based on existing literature we hypothesized that the type of immunosuppressant had a distinct effect on GM composition. Results were therefore stratified based on the primary immunosuppressant, namely TAC or CSA. There was no difference in total bacterial cell count and BC to HC between the two groups (**Figure 3A**). Patients receiving TAC as the primary immunosuppressant had a significant lower Shannon diversity (3.36 ± 0.47) compared with both HC (3.89 ± 0.39) and with patients receiving CSA (3.70 ± 0.30). Similarly, the number of observed species was significantly lower in the TAC group (85.2 ± 40.7) compared with HC (127 ± 34.8) and the CSA group (121 ± 26.7). Differences to the CSA group remained significant when organ was included as a random effect. No difference in diversity between patients receiving CSA and HC were found. Moreover, we observed a positive correlation between the TAC concentration in blood and the BC to HC dissimilarity across all patients (**Figure 3B**); this observation remained when organ was included as random effect, albeit only as a trend (*p=0.0591*). Increased TAC levels were also associated with lower diversity in LTR as observed by a negative correlation between TAC blood levels and the number of observed species; this relation was specific for LTR and not observed when samples from HTR and KTR recipients receiving TAC were included, even when organ was included as random effect. The CSA levels did not correlate with any of the above analyzed variables.

**Figure 3.**
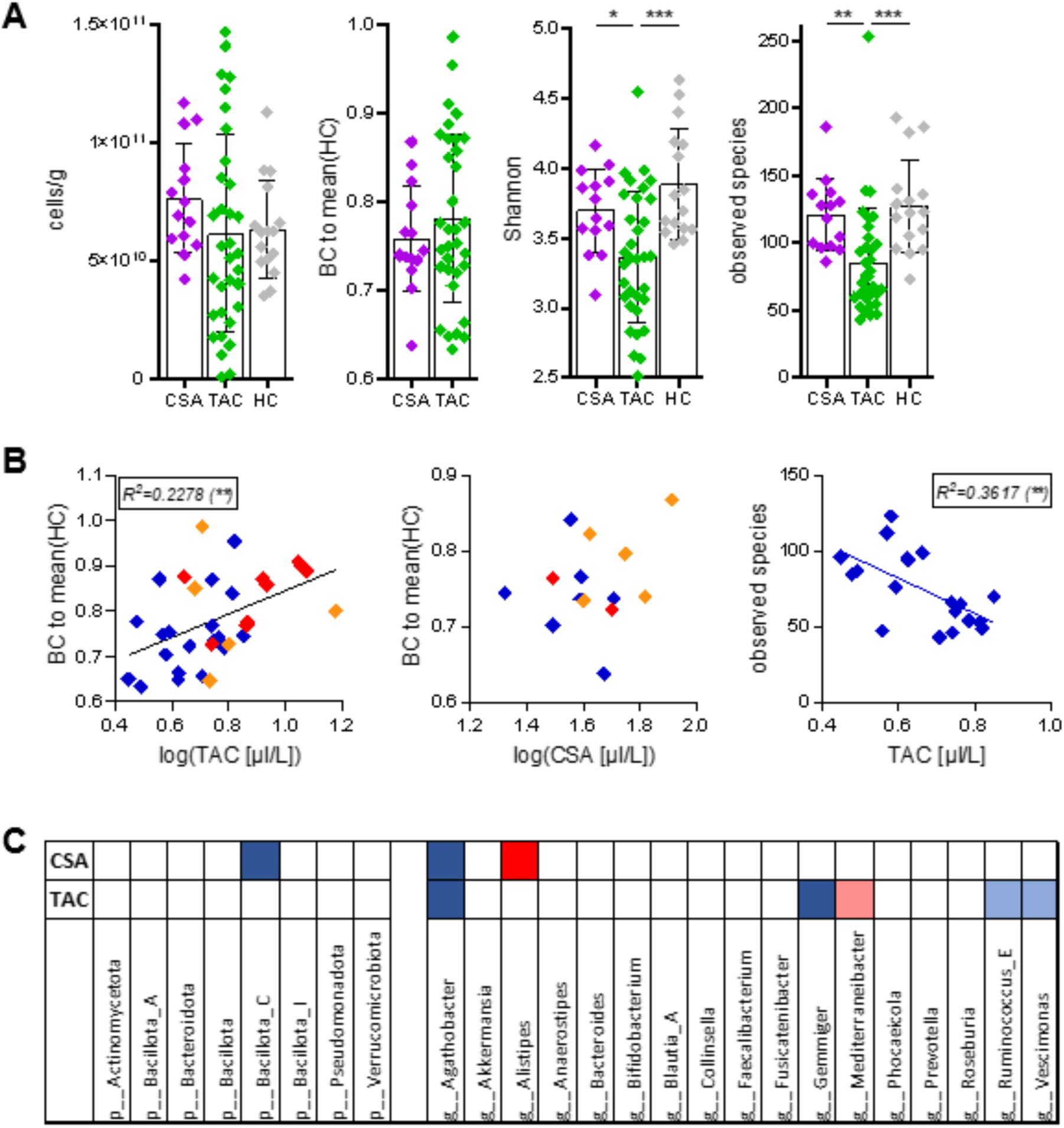
Comparison of samples based on the primary immunosuppressant administered. The distribution of bacterial cell count, Bray Curtis dissimilarities (BC) to healthy controls (HC), the Shannon index and the number of observed species per immunosuppressant group and HC is depicted in panel **A**. In panel **B** the association of BC to HC and tacrolimus (TAC) levels (left) or Cyclosporine A (CSA) levels (center) are visualized. The right panel shows the association between TAC levels and the number of observed species in samples from liver transplant recipients. Models were calculated based on linear regression. The table in panel **C** gives information about significant differences of the most abundant phyla (left) and genera (right) for the TAC and CSA group compared with HC (for color scheme see Figure 2). * - p<0.05, ** - p<0.01, *** - p<0.001, **** - p<0.0001.

Investigating GM composition showed a decreased relative abundance of Agathobacter and Gemmiger in patients receiving TAC as the primary immunosuppressant compared with HC. Furthermore there was a trend towards lower relative abundances of Ruminococcus_E and Vescimonas and a higher relative abundance of Mediterraneibacter. For the CSA group, a decreased abundance of bacteria belonging to the phylum Bacillota_C compared with HC was found. On the genus level, there was a significant higher relative abundance of Alistipes and a lower relative abundance of Agathobacter in the CSA group compared with HC (**Figure 3C**).

### C2 showed an increased SIgA concentration and targeting

The specific focus of this study was to investigate whether immunosuppression affected the fecal SIgA concentration and its GM targeting. The percentage of SIgA coated bacteria ranged from 1.3 % to 78.9% across all samples and, overall, patients trended towards a higher coating of GM by SIgA (22.6 ± 17.3%) compared with HC (12.9 ± 6.5 %, *p=0.0546*) (**Figure 4A**). On the cluster level, C2 had a significantly higher proportion of SIgA coated GM (30.2 ± 19.6 %) compared with both C1 (14.7 ± 9.8 %) and HC (**Figure 4A**). That was accompanied by a significantly higher total fecal SIgA concentration (1139 ± 1569 µg/g) compared with HC (445 ± 685 µg/g) and C1 (685 ± 1969 µg/g) (**Figure 4B**). On the organ level, HTR had a higher proportion of SIgA coated GM (32.8 ± 17.1 %) compared with both HC and KTR (14.6 ± 5.4 %) (**Figure 4A**). LTR and KTR did not differ in their proportion of SIgA coated GM compared with HC.

**Figure 4.**
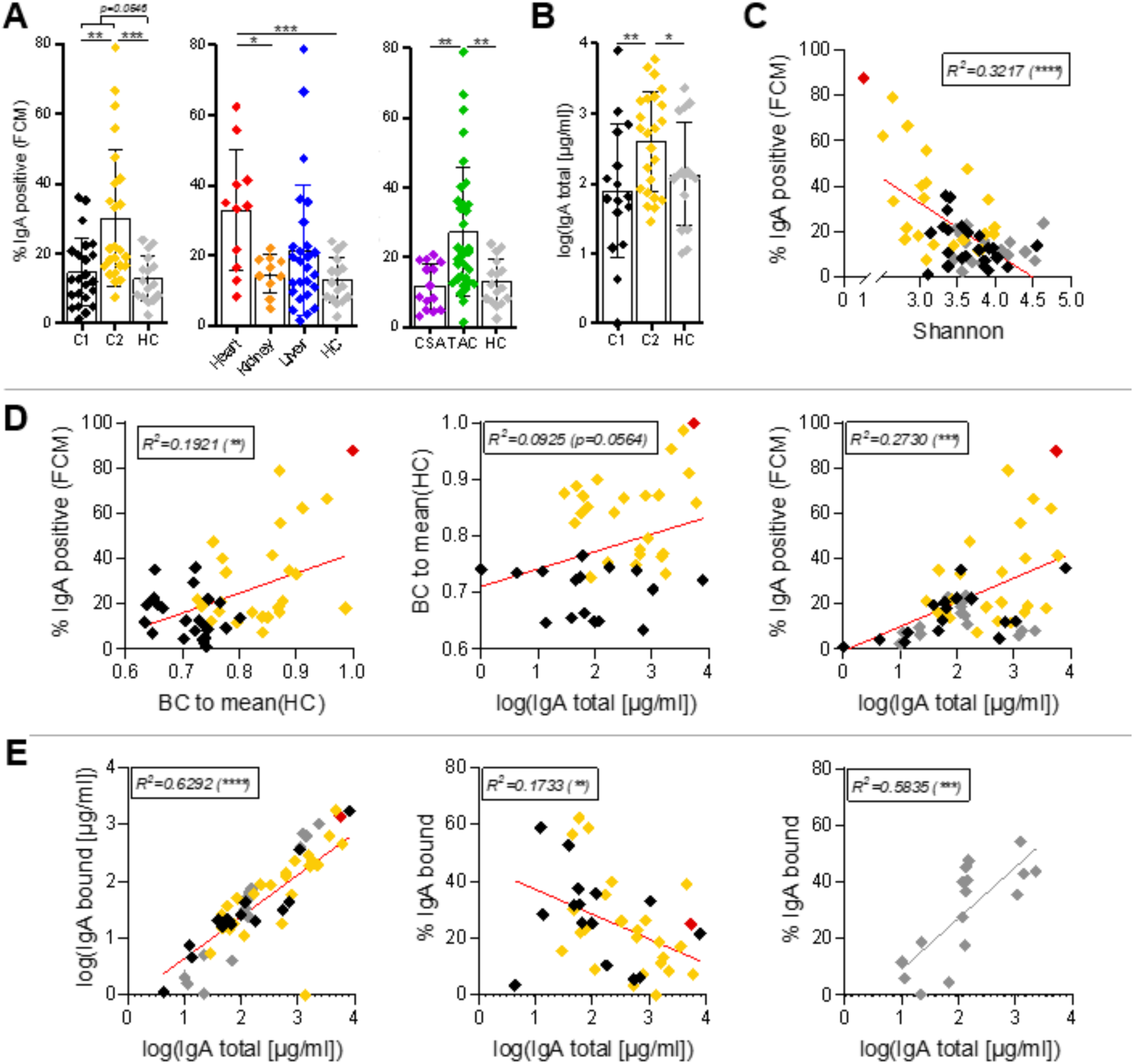
The proportion of SIgA coated bacteria (panel **A**) is compared between clusters (left), organ (center) and primary immunosuppressant (right). In panel **B** the total SIgA concentration of clusters and healthy controls (HC) is shown. In panel **C** the association of the proportion of SIgA coated bacteria and the Shannon index is depicted whereas in panel **D** the associations between the proportion of SIgA coated bacteria, total SIgA concentration and Bray Curtis dissimilarities (BC) to HC are compared across all samples. Panel **E** gives the association between total SIgA concentration as well as the concentration (left) and percentage (center, right) of bound SIgA, respectively. Models in panels **C-E** were calculated based on linear regression. Colors indicate the allocation to the respective cluster (black = C1, gold = C2, grey = HC). The sample L07 (excluded from analysis) is indicated in each graph as a red diamond.* - p<0.05, ** - p<0.01, *** - p<0.001, **** - p<0.0001.

Analyses based on the primary immunosuppressants revealed a significantly higher percentage of GM coated by SIgA in patients receiving TAC (27.2 ± 18.4 %) compared with patients receiving CSA (11.9 ± 3.9 %) as well as with HC (**Figure 4A**); no difference in the relative SIgA coating was found for the CSA group when compared with HC. For the sample dominated by Klebsiella (L07) the majority of bacteria (87.5 %) were coated by SIgA (**Supplementary Figure S3**).

A lower Shannon Index was associated with a higher proportion of SIgA coated GM in patients (*p<0.0001*), however, not in HC (*p=0.7432*) (**Figure 4C)**. Dysbiosis (BC to HC) was positively associated with the proportion of SIgA coated GM and the total SIgA concentration in patients, albeit the latter only as a trend (*p=0.0564*) (**Figure 4D**). Moreover, an increase in total SIgA concentration was positively associated with an increase in the proportion of SIgA coated GM in patients (**Figure 4D**). Furthermore, there was a strong relation between the total SIgA concentration and the concentration of bound SIgA (**Figure 4E**) indicating that increasing SIgA secretion is indeed associated with increased targeting of GM. However, the total SIgA concentration was negatively associated with the proportion of bound SIgA in patients suggesting that, despite an increased SIgA binding with higher levels of total SIgA, a relative larger fraction remained not bound to bacteria (**Figure 4E**); this was not found for HC, where increased total SIgA concentration was connected with higher percentages of bound SIgA.

### Identification of specific taxa targeted by SIgA

We characterized GM community within SIgA positive and negative fractions after cell sorting based on 16S rRNA gene sequencing and compared results with total GM composition to investigate SIgA target spectra. For both clusters and HC we observed a significant separation of the whole community (star) from both the SIgA positive fraction (dot) and the SIgA negative fraction (triangle) based on PERMANOVA analyses demonstrating specific targeting of GM by SIgA (**Supplementary Figure S4**). On the organ level, LTR showed, similar as HC, a significant separation of the total community from both fractions based on PERMANOVA analyses. For KTR, the fraction targeted by SIgA was significantly different from total GM composition (no difference with the negative fraction was observed), whereas no overall differences between total GM composition and any fraction were seen in HTR due to highly subject-specific patterns of communities in those patients.

To identify individual taxa that showed a shift in SIgA-targeting, we calculated the SIgA-binding Probability (SIgA-bP) as described in Jackson and colleagues^26^ under the prerequisite that a respective taxon was present in at least five samples per individual group analyzed. Considering the abundance of the respective taxon in each group, we first identified taxa that were overall strongly or hardly targeted by SIgA, as defined by an average SIgA-bP of greater than 0.25 or lower than -0.25, respectively, over all samples, which resembles a >50-fold higher abundance in one of the fractions. Three taxa were identified that were highly targeted across all samples, namely Eubacterium_I (SIgA-bP = 0.35 ± 0.45, n=23), Limivivens (0.36 ± 0.42, n=12) and Scatomorpha (0.31 ± 0.42, n=15) (**Figure 5A**). Another seven taxa had a strong negative SIgA-bP across all samples, namely Anaerobutyricum (-0.29 ± 0.40, n=23), Coprococcus (-0.30 ± 0.37, n=15), Gallintestinimicrobium (-0.40 ± 0.43, n=23), Hominicoprocola (-0.28 ± 0.36, n=14), Hominilimicola (-0.36 ± 0.45, n=32), Phascolarctobacterium (-0.29 ± 0.44, n=13) and Roseburia (-0.40 ± 0.34, n=31). All identified taxa with an overall large (negative) SIgA-bP had a mean relative abundance below 1 % with the exception of Roseburia (**Supplementary Figure S5**).

**Figure 5.**
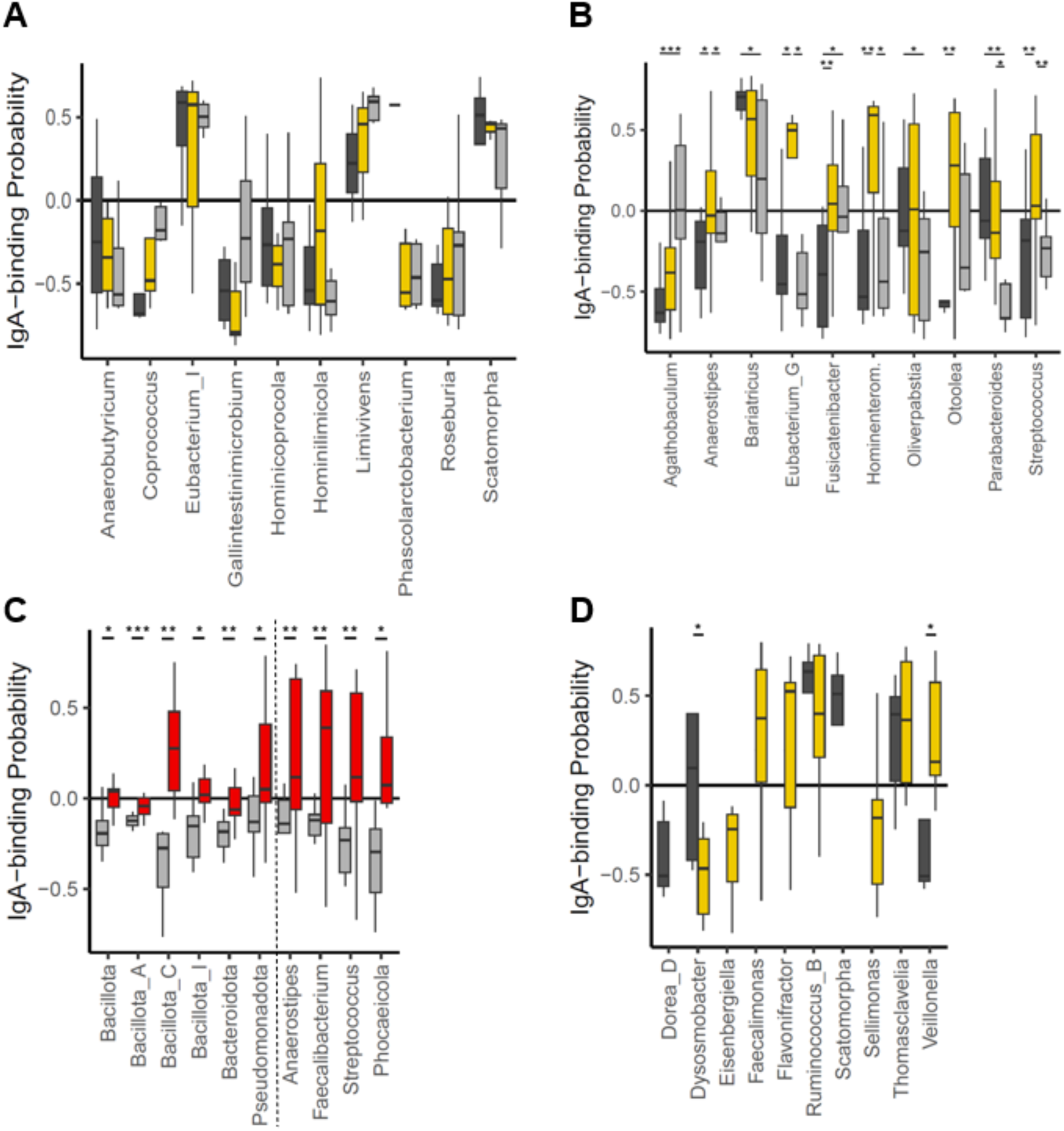
Comparison of SIgA binding Probabilities (SIgA-bP) of taxa between groups. Panel **A** depicts taxa with a mean SIgA-bP >0.5 or <-0.5 across all samples. SIgA-bP distribution is given per cluster and healthy controls (HC). Taxa with significant different SIgA-bP between clusters and HC and between heart transplant recipients (HTR) and HC are shown in panel **B** in panel **C**, respectively. The dotted line (panel **C**) separates phyla from genera. Panel **D** depicts the SIgA-bP of taxa that were strongly targeted in clusters but were not abundant in HC. Colors indicate group allocation (dark grey = C1, gold = C2, red = HTR, light grey = HC). * - p<0.05, ** - p<0.01, *** - p<0.001, **** - p<0.0001.

Next, we focused on taxa that were differently targeted between clusters, as defined by a significant difference in SIgA-bP. Analysis of SIgA targeting patterns was focused on the cluster level as previous analyses revealed this level provided clear separation between patients and comprised high sample numbers per group. On the phylum level, only bacteria from Bacillota_A were significantly more targeted by SIgA in C2 compared with both HC and C1 (**Supplementary Figure S5A**). Additionally, there was a trend towards a higher SIgA targeting of bacteria belonging to the phyla Bacillota_I (*p=0.0512*) and Pseudomonadota (*p=0.0561*) in C2 compared with HC. On the genus level, Anaerostipes was significantly more targeted in C2 and Fusicatenibacter was significantly less targeted in C1 compared with both HC and the respective other cluster (**Figure 5B**). Additionally, eight taxa with a mean relative abundance <1% were identified as differentially targeted between clusters and HC (**Figure 5B**). Bacteria belonging to the genera Agathobaculum and Parabacteroides were significantly less and more targeted, respectively, in all patients compared with HC (Figure 5B). On the cluster level, samples from C1 had a higher SIgA-bP of Bariatricus and Oliverpabstia compared with HC. Eubacterium_G, Hominienteromicrobium and Streptococcus were significantly more targeted by SIgA in C2 compared with HC and C1. The taxon Otoolea showed significantly different targeting between clusters, being relatively more targeted in C2 compared with C1 (**Figure 5B**).

For providing comprehensive analyses, we also investigated differences in SIgA-bP on the immunosuppressant and organ levels. On the immunosuppressant level, Bacillota_A was significantly more targeted by SIgA in TAC compared with both HC and with patients receiving CSA. On the genus level, Parabacteroides was more targeted by SIgA whereas Agathobaculum was less targeted by SIgA patients receiving TAC compared with HC. Patients receiving CSA showed an increased SIgA-bP of Hominilimicola and a decreased SIgA-bP of Hominimerdicola compared with HC (data not shown); no differences in SIgA-bP between the TAC and CSA group on the genus level were found. On the organ level, HTR showed the most unique SIgA targeting patterns, which included several taxa that were not identified on any other level. On the phylum level there were strong increases in SIgA-bP, where almost all phyla with a mean relative abundance >0.1 % were more targeted by SIgA in HTR compared with HC (**Figure 5C**). On the genus level, four genera, namely Anaerostipes, Faecalibacterium, Phocaeicola and Streptococcus, were significantly higher targeted by SIgA in that group compared with HC (**Figure 5C**). In LTR and KTR Agathobaculum had a significantly lower SIgA-bP and Parabacteroides had a significantly higher SIgA-bP compared with HC, respectively (data not shown).

Finally, we investigated taxa that were specifically associated with patients’ GM and therefor contributed to the observed dysbiosis in SOT recipients. We focused on taxa that were only present in patients or a specific patient cluster and largely devoid in HC and selected those that were overall strongly or hardly targeted by SIgA as defined before (SIgA-bP of +/-0.25). The taxa Ruminococcus_B and Thomasclavelia were present in the majority of patient samples (n=32 and n=30, respectively) and were mostly absent in HC. Both taxa were highly targeted by SIgA with an average SIgA-bP of 0.42 ± 0.38 and 0.32 ± 0.32, respectively (**Figure 5D**). Bacteria belonging to the taxa Dysosmobacter and Veillonella were only present in patients and significantly differed in their SIgA-bP between clusters, the former being more targeted by SIgA in C1, the latter in C2 (**Figure 5D**). Three taxa were predominantly present in C2 and showed strong SIgA-bP values, namely Eisenbergiella, Flavonifractor and Sellimonas, whereas Faecalimonas had a strong negative SIgA-bP in C2. C1 was characterized by a strong negative SIgA-bP of Dorea_D and a strong positive SIgA-bP of Scatomorpha, both of which were hardly abundant in C2 (**Figure 5D**).

## Discussion

Life-long immunosuppression after SOT is crucial to prevent the rejection of the transplanted organ. However, it is also associated with alterations in the GM composition.^8^ We confirmed persisting GM dysbiosis in SOT recipients even several years post transplantation and showed that the extent of GM dysbiosis was associated with the overall immunosuppressant regimen, in particular with TAC levels. Additionally, we demonstrated that the total SIgA concentration and the proportion of SIgA coated bacteria are elevated in patients, which was associated with the degree of dysbiosis and altered SIgA target spectra.

The observed GM dysbiosis in our cohort is in accordance to existing literature investigating GM composition in human adult and pediatric SOT recipients.^14,27^ Identifying and targeting factors contributing to GM dysbiosis is crucial for the short- and long-term health of SOT recipients. In short-term post-surgery GM dysbiosis is associated with several complications including an increased risk for infection^12^, diarrhea^13,14^ and acute rejection (as reviewed in ^10,15^). Regarding the long-term development, we recently demonstrated an inverse correlation between GM diversity and transaminase levels as indicator for graft health and were further able to predict hepatocellular damage of the transplant based on GM with high accuracy.^2^ A recent large-scale analysis of 1337 metagenomes from a longitudinal follow-up cohort by Swarte et al.^11^ further highlighted the relevance of targeting GM dysbiosis in SOT recipients as it was strongly positively associated with long-term all-cause mortality as well as cause-specific mortality in those patients.

We collected samples at least one year post transplantation to exclude any direct influences on GM from surgery and associated short-term treatment thereafter and further excluded any patients with recent antibiotic intake enabling the hypothesis that GM dysbiosis in our cohort was indeed mainly caused by drug-mediated effects. We observed a strong and dose-dependent effect of TAC on GM dysbiosis that was not observed in patients receiving CSA as the primary immunosuppressant. Furthermore, the multidrug-regimen in HTR and KTR might have contributed to the higher GM dysbiosis in those patients compared with LTR. The association between drug-induced immunosuppression and GM dysbiosis was already observed in earlier studies based on both animal models and humans.^28–30^ However, as most studies were observational, the question of the underlying mechanism by which immunosuppressants affect GM composition remains open. One potential mechanism is the direct antimicrobial effect of immunosuppressants on GM. A large-scale study from Maier and colleagues^31^ in 2018 screened more than 1000 non-antibiotic drugs on their impact on selected GM strains including also some immunosuppressants, namely CSA, prednisolone and azathioprine. Only CSA and azathioprine exerted an antimicrobial effect acting on five and seven of 40 total strains, respectively, whereas no antimicrobial effect was observed for prednisolone; authors did not investigate TAC, which was the main immunosuppressant in this study. Another route for drugs altering GM composition might be via the immune system, which was the focus of only a few studies. For instance, Tourret and colleagues^32^ found that altered GM composition was associated with reduced expression of C-type lectins and IL-22 after 14-days upon TAC treatment. In general, disentangling direct effects of immunosuppressants on GM from effects mediated via alterations of the immune system is a difficult task and remains largely elusive. This problem was also pointed out in a systematic review from Manes and colleagues comparing existing studies to investigate the bidirectional interaction between immunosuppressants and GM.^29^ The authors highlight that results are largely based on observational studies and comparison of results is heavily confounded by heterogeneity of administered drug(-combinations), inclusion and exclusion criteria, the overall study design as well as the GM modulating effect of the underlying disease itself. Nevertheless, the authors concluded that immunosuppressants exert their antimicrobial effect most likely via modulating the host immune response rather than via a direct interaction with GM.^29^

The initial hypothesis of this study was that immunosuppression results in a reduced IgA production and GM targeting as it acts on T and B cell proliferation and differentiation that are important for IgA production. However, the obtained results contradicted this hypothesis. In contrast, we observed higher SIgA concentration and GM targeting in SOT patients, which were further associated with higher immunosuppressant levels as well as with increased numbers of different types of immunosuppressants. We created a hypothetical model connecting immunosuppression, GM composition and SIgA targeting in order to integrate our results in a larger mechanistical framework based on existing literature (**Figure 6**). First, we observed that the extent of GM dysbiosis in SOT recipients was associated with increased immunosuppression levels, at least with TAC, which is in line with previous studies as discussed above. Furthermore, GM dysbiosis in patients was positively associated with increased and altered targeting of GM by SIgA, which was additionally connected with increased SIgA concentrations. Observations from literature suggest that higher SIgA levels in patients are routed in a regulatory T cell (T_reg_) mediated activation of IgA+ B cells by immunosuppressants, and we hypothesize that this activation promotes an increased SIgA-mediated host immune response towards GM. Taken together, our model suggests that both direct drug-related effects on GM and via the immunes system, in particular via SIgA, promote dysbiosis in SOT patients.

**Figure 6.**
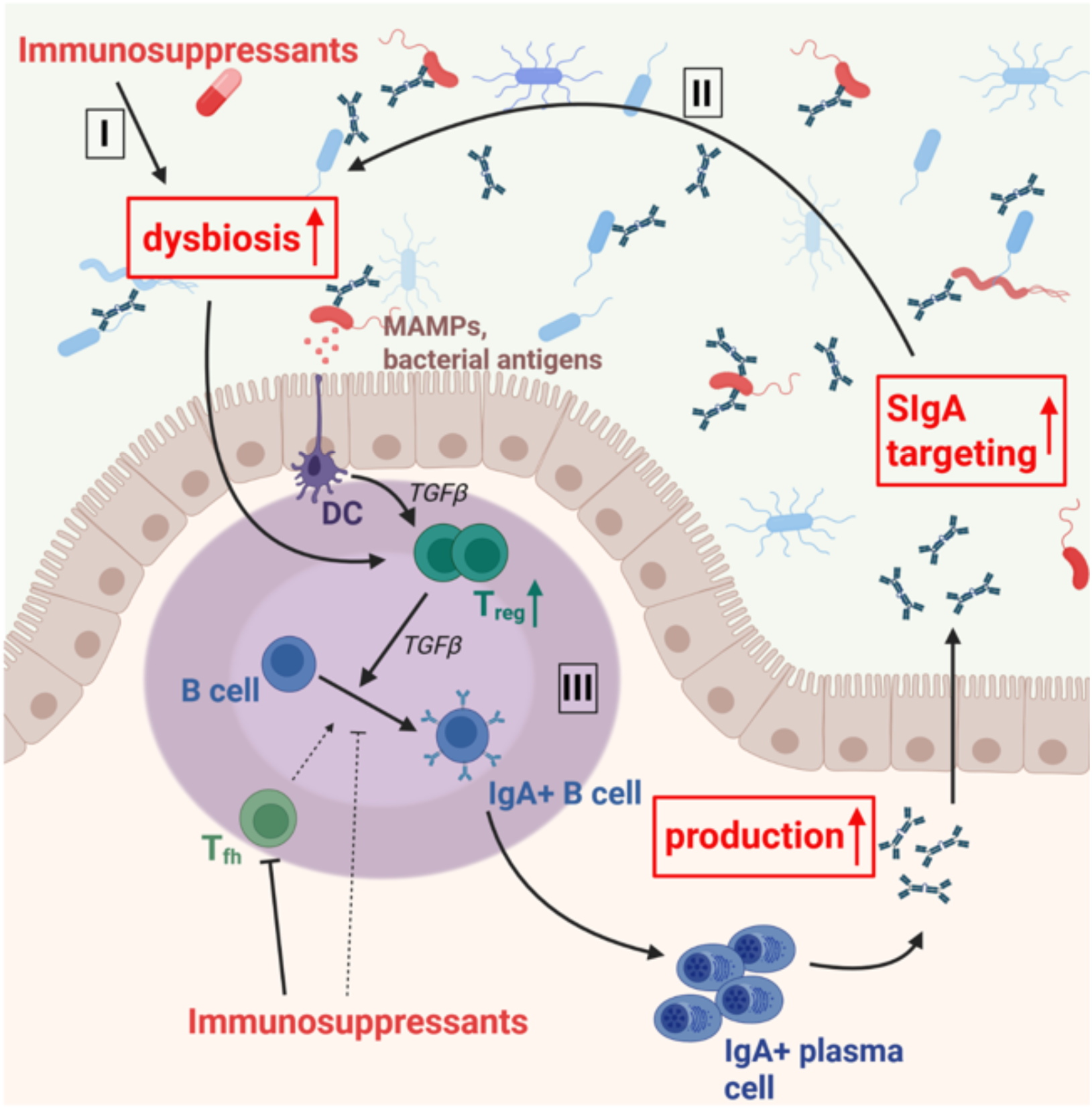
Proposed model connecting immunosuppression, gut microbiota (GM) composition and SIgA targeting. Drug-induced immunosuppression **(I)** together with increased SIgA concentration and targeting **(II)** are contributing to dysbiosis in SOT recipients. Higher SIgA production is mediated via the activity of regulatory T cells (T_reg_) that react to signals from dysbiosed GM **(III)**.

GM dysbiosis was positively associated with increased SIgA targeting in our cohort and especially pronounced in patients allocated to C2. Patients in this cluster were characterized by an overall increase in SIgA concentration and proportion of SIgA coated bacteria as well as specific alterations in the SIgA targeting pattern. The association between higher SIgA levels and GM dysbiosis was already observed in other disease cohorts, e.g. inflammatory bowel disease (IBD)^33,34^, psoriatric arthritis^35^ and schizophrenia.^36^ The potential consequences of increased SIgA targeting of GM increasing dysbiosis remain open. On the one hand, SIgA targeting is generally regarded as host-mediated mechanisms for pathogen exclusion and promotion of a eubiotic GM community in a healthy system. Consequently, one can hypothesize that the host immune system increases SIgA production to resolve drug-related GM dysbiosis. Based on this hypothesis, we would have observed higher targeting of pathogens and GM members that showed increased relative abundances in patients. However, our results suggest increased SIgA targeting of commensal GM that showed no differences in relative abundances in patients compared with HC. Furthermore, bacteria belonging to genera that are associated with pathogenic features, such as Escherichia or Hungatella, were not more targeted by SIgA in patients. Taken together those observations weaken the hypothesis that altered SIgA targeting of GM in patients is a specific response of the host immune system to conquer GM dysbiosis. Rather, results indicate that altered SIgA production and target spectra are the consequence of a dysfunctional immune system and, hence, contribute to dysbiosis by creating growth niches for more pathogenic bacteria and reducing the growth environment of commensals. We, hence, decided to include altered SIgA response in SOT patients as a dysbiosis-promoting factor in our model presented in **Figure 6**.

As discussed, we found higher SIgA production despite an expected immunosuppressant-mediated suppression of T and B cell activation and proliferation.^37–40^ Based on literature we propose a dose-dependent local increase in regulatory T cells (T_reg_) that leads to the stimulation of B cell proliferation and class switch recombination (CSR) resulting in increased IgA production. The association between TAC administration, GM dysbiosis and increased T_reg_ numbers was observed in several animal studies that hypothesized that T_reg_ production is rather mediated via GM than by direct effects of the immunosuppressant itself.^30,41^ Other studies further highlighted the importance of T_reg_ for B cell CSR that is signaled via transforming growth factor β (TGFβ).^42,43^ Aside from TAC, also prednisolone was associated with higher T_reg_ numbers both *in vitro* and in KTR comparing the numbers before and four month after transplantation.^44,45^ Evidence regarding the effect of CSA is scarce but point to an inhibitory effect on T_reg_. ^46,47^ This further supports a T_reg_ mediated increase in SIgA production in our cohort as we did not see increased an SIgA concentration in patients receiving CSA as the only immunosuppressant. Taken together, we postulate that continuous intake of immunosuppressants is causing GM dysbiosis and subsequently stimulates SIgA production via T_reg_. The increased and altered SIgA-targeting of GM further enhances GM dysbiosis thus creating a positive feedback loop where SIgA-mediated dysbiosis is fueling increased SIgA production along with alterations in target spectra.

This study has some limitations due to its design as mono-centric study limiting the total number of patients and resulting in a lower sample size. Furthermore, the heterogeneity of immunosuppressant-regimens was especially high in KTR and HTR that might have masked effects from individual primary immunosuppressants. Higher sample numbers would allow a more in-depth analysis of the contribution of each immunosuppressant to the observed effects. We only collected one sample per patient performing cross-sectional analyses across organs and immunosuppressant groups. While it allowed for uncovering major signals, a longitudinal approach would provide the basis for more detailed analyses on SIgA responses in the context of GM dysbiosis development.

In summary, while immunosuppression is administered as a life-long therapy to prevent graft rejection, it is also associated with a dysbiosed GM composition that remains stable even years after transplantation, thereby increasing the risk for (long-term) complications. Our study highlights the crucial role of SIgA at the interface between the host immune system and GM in SOT recipients and suggests a contributing role of SIgA for the observed GM dysbiosis in those patients. The results give rise to SIgA as a potential novel target to combat dysbiosis in SOT recipients.

## Methods

### Study cohort and sample collection

All patients were recruited at the pediatric hospital at Hannover Medical School. Inclusion criteria were the transplantation of a solid organ (heart, kidney or liver) with the time of transplantation >1 year. Exclusion criteria were antibiotic therapy within one month before sampling and any diseases affecting gastrointestinal health, bile acid metabolism or pancreatic secretion. Fecal samples were collected at home or during a hospital visit and brought to the hospital upon the next visit. A naïve aliquot and an aliquot stored in RNA/DNA shield (ZYMO, USA) were collected by the participants and stored at -20°C until processing.

### Processing of fecal samples for flow cytometric (FCM) analyses

Bacteria (cells per gram stool) were measured as described in Kircher et al.^48^ Subsequently three aliquots containing 50 million bacteria each were taken for SIgA processing, one aliquot served for quantification of SIgA targeted bacteria, one aliquot was used for cell sorting and the third aliquot served as an (unlabeled) control. All aliquots were centrifuged at 8,000 x g at 4°C for 5 minutes and the supernatant was discarded. Samples were washed with cold staining buffer (1xPBS, 1% BSA, 2mM EDTA) and the supernatant was again discarded. The bacterial pellet was blocked for 20 minutes on ice using mouse serum (1:10) before incubating with APC-conjugated anti-human IgA (1:10, Miltenyi, Germany) in the case of the first two aliquots and with 1xPBS (1:10) in the case of the control aliquot for 30 minutes covered on ice. Following incubation, cells were again washed twice with cold staining buffer, and the supernatant was discarded. Bacterial pellets of the cell sorting aliquot and the control aliquot were resuspended in 200 µl staining buffer whereas the remaining bacterial pellet was resuspended in 1ml PBS for flow cytometric SIgA quantification. All samples were incubated with SYBR Green I (Thermo Fisher Scientific, USA) and 0.5 mM EDTA for 15 minutes at RT. The aliquot for sorting was stored on ice and sorted within one hour whereas the aliquot for FCM was immediately measured at a Northern Lights (Cytek, USA). The percentage of SIgA coated bacteria was quantified by plotting the signals detected from SYBR Green I emission (all bacteria) against the signal detected from the APC-conjugated anti-IgA (SIgA-coated bacteria). The measurements from the control sample served to set the gate accordingly. An examples for the gating is shown in Supplementary figure 3.

### Fluorescent activated cell sorting of bacteria

Bacterial populations were sorted into an IgA-positive fraction, containing cells that were bound by the anti-IgA antibody, and an IgA-negative fraction, with cells that were not targeted. 500,000 cells were sorted for each fraction using a FACSAria Ilu (BD, USA) with a 70 µm nozzle. After sorting, both fractions were remeasured and resorted if the purity was below 80 %.

### Sequencing and bioinformatics analyses

DNA was extracted using the ZymoBIOMICS Miniprep Kit (ZYMO, USA) for the naïve samples and the ZymoBIOMICS Mikroprep Kit (ZYMO, USA) for the FACS fractions according to the manufacturer’s protocols. Library preparation for 16S rRNA gene sequencing (IgA fractions and naïve samples) and metagenomics shotgun sequencing (only naïve samples) and subsequent sequencing on MiSeq and NovaSeq6000, respectively, was done as described in Kircher et al.^48^ Bioinformatic analysis of amplicons (16S rRNA gene) was performed as described in reference ^48^, whereas analyses of shotgun data was done according to reference ^49^ yielding compositional and functional data.

### Measurement of SIgA fecal concentration

Total fecal SIgA concentration and the proportion of bound and unbound SIgA were quantified using both the whole filtrate and the supernatant of 1:10 diluted naïve fecal samples in PBS based on an IgA uncoated human Enzyme-Linked Immunosorbent Assay (ELISA, Thermo Fisher Scientific, USA) according to the manufacturer’s instruction. Concentration in the filtrate was indicating total fecal SIgA whereas the concentration in the supernatant was attributed as unbound SIgA. The concentration of bound SIgA was calculated by subtracting the unbound SIgA values from those of the total SIgA.

## Statistical analysis

Hierarchical clustering was done by k-means analyses based on Bray Curtis (BC) dissimilarities on the species level using R Statistical Software (version 4.3.3, R foundation for statistical Computing, Vienna, Austria) with the *cluster* (v2.1.6) and *purrr* package (v1.0.2). Metric dimensional scaling analysis was done on BC dissimilarities on the species level for metagenomics data and on the genus level for 16S rRNA sequencing data using the *phyloseq* package (v1.44.0). PERMANOVA analyses were performed with the *adonis* function of the *vegan* package (v2.5.7) on BC dissimilarities (genus level). GM diversity was compared between groups via calculating the Shannon Index, the number of observed species, and the BC distance to the average BC dissimilarity in HC (BC to HC) for each sample. For differential abundance calculations of taxa, the relative abundance of a taxon was compared to the mean relative abundance of the respective taxon within HC using linear regression analyses (function *lm*; log(data +1)) and corrected for multiple testing using a local false discovery rate (lfdr) <0.05, calculated via the *fdrtool* package (v1.2.18). An lfdr <0.1 was considered as a trend. General characteristics of patients, diversity and SIgA parameters were compared between groups using the *lm* function of the package fdrtool (v1.2.18) feeding log-transformed data (log(data+1)) and controlling for the time since transplantation. For all parameter a *p*-value < 0.05 was considered significant. On the immunosuppressant level, organ was included as random effect for comparison between parameters using the *lmer* function of the package lme4 (v1.1.36). Selected parameters were correlated in R using regression analyses (function *lm*) feeding log-transformed data (log(data +1) for continuous variables. Boxplots and scatter plots were created with GraphPad Prism (version 8.4.3). The SIgA binding Probability was calculated according to the equation given in reference ^26^.

### Ethics

Our study was approved by the ethics committee of Hannover Medical School and respected the declarations of Helsinki. All patients gave permission for the analysis of their data and biomaterial in the form of written informed consent.

## Acknowledgements

We thank all participants of the study for providing sample.

---

**Supplementary figure 1.**
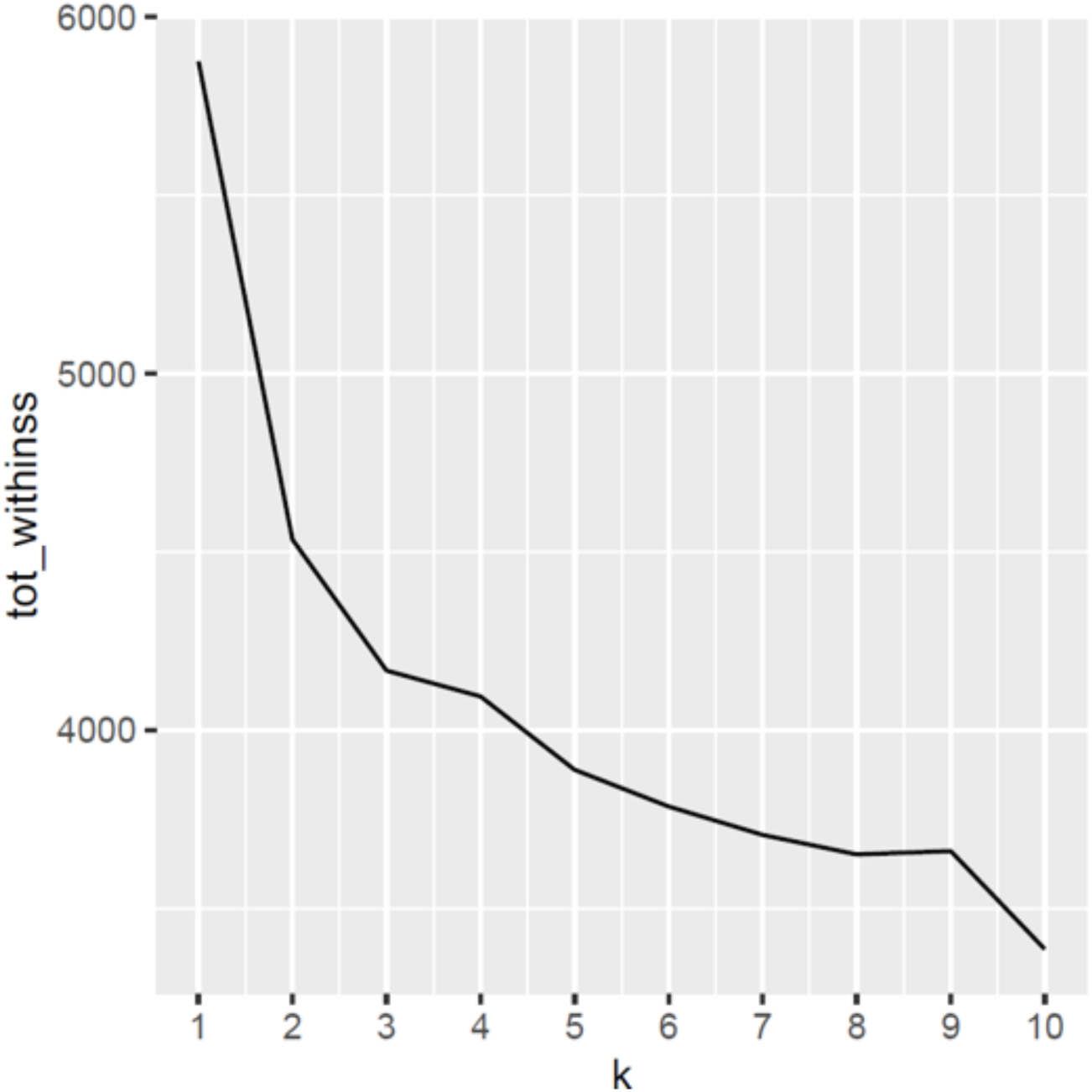
Elbow plot from k-means based clustering algorithm to identify the optimal number of clusters using metagenomics data on species level and after the exclusion of sample L07. According to this analysis a k of 2 is optimal to stratify samples.

**Supplementary figure 2.**
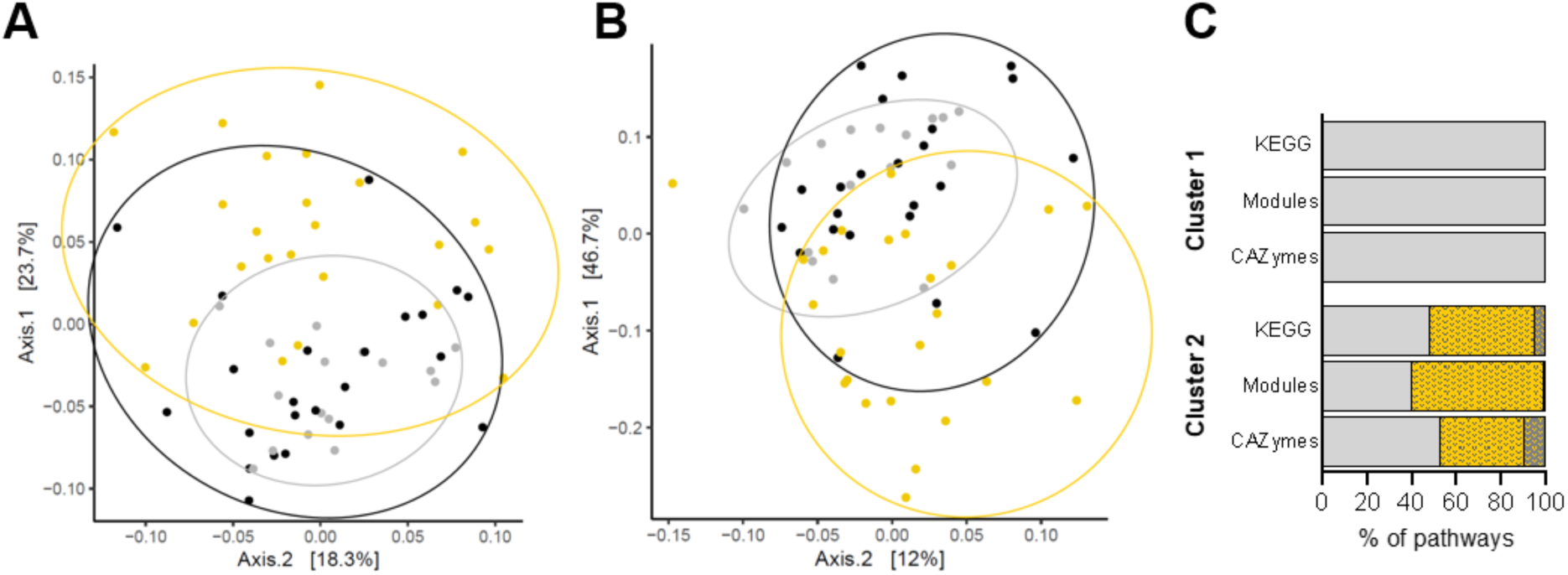
Functional analysis of bacterial communities based on metagenomics data. Metric multidimensional scaling analysis of all samples based on Bray Curtis dissimilarities of the abundances of KEGG orthologues (panel **A**) and CAZymes (panel **B**). Samples are colored based on the allocation to the identified clusters from the taxonomy-based analysis shown in figure 1A and the respective cluster distribution was further visualized by circles. Panel **C** indicates the percentage of pathways with a similar (light grey), higher (gold) or lower (dark grey) abundance in patient clusters compared with healthy controls (HC).

**Supplementary figure 3.**
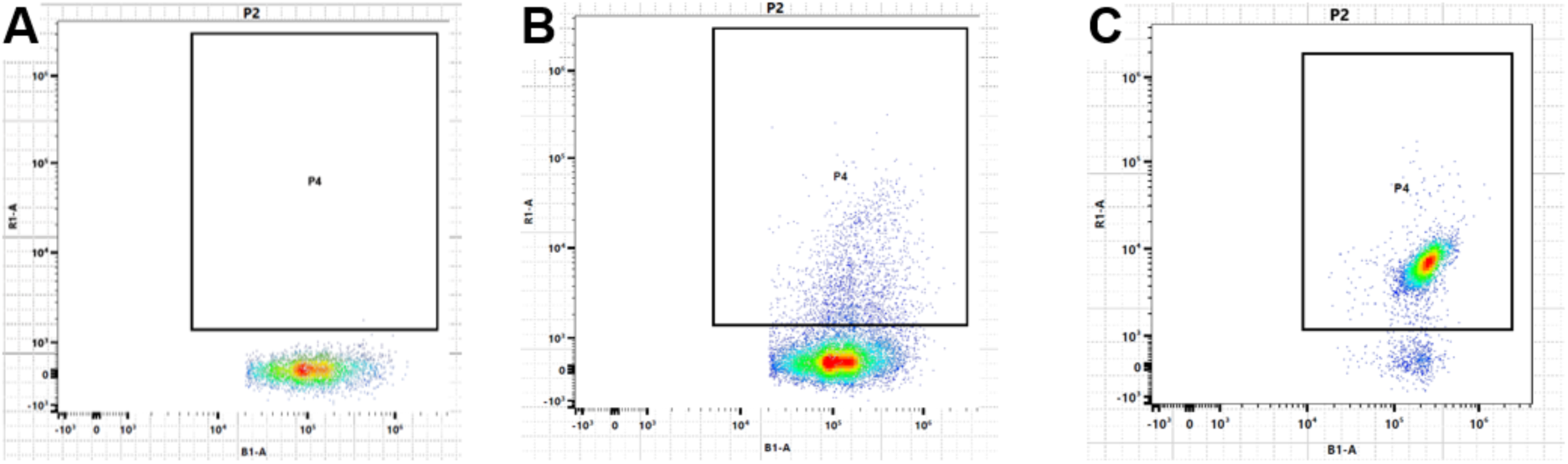
Exemplary images of flow cytometric analysis of samples before (panel **A**) and after (panel **B**) incubation with the anti-IgA antibody. Gate P4 included all bacteria that were bound by the anti-IgA antibody. For sample L07, most bacteria (87.5 %) were SIgA positive (panel C).

**Supplementary figure 4.**
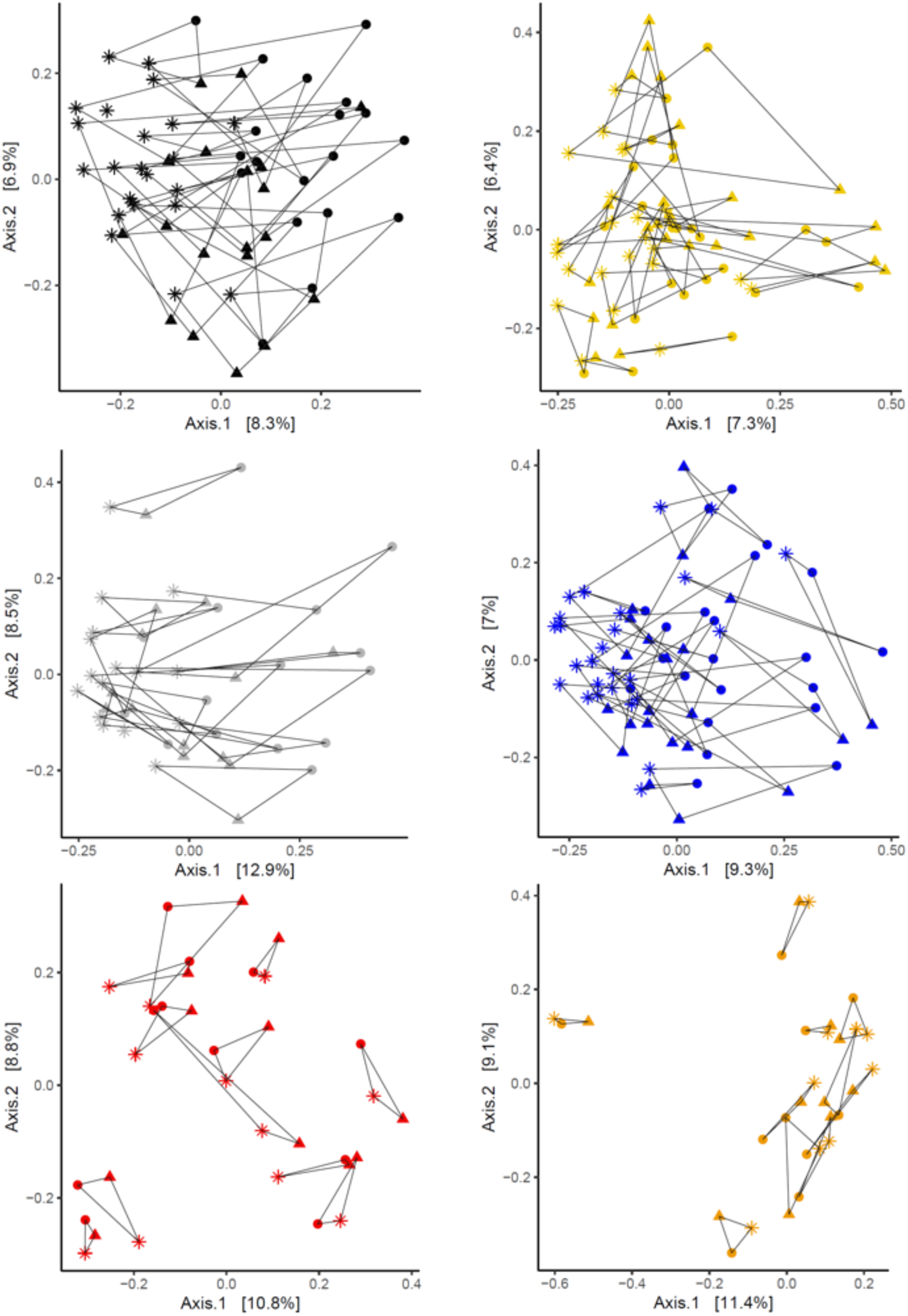
Metric multidimensional scaling analysis based on Bray Curtis dissimilarities of 16S rRNA gene sequencing data comparing bacterial compositions of the unsorted samples (star), the positive fractions (dot) and the negative fractions (triangle) per cluster (upper panels) and per transplanted organ (lower panels). Lines are connecting sample of each subject. Samples are color coded according to groupings shown in Figure 1.

**Supplementary figure 5.**
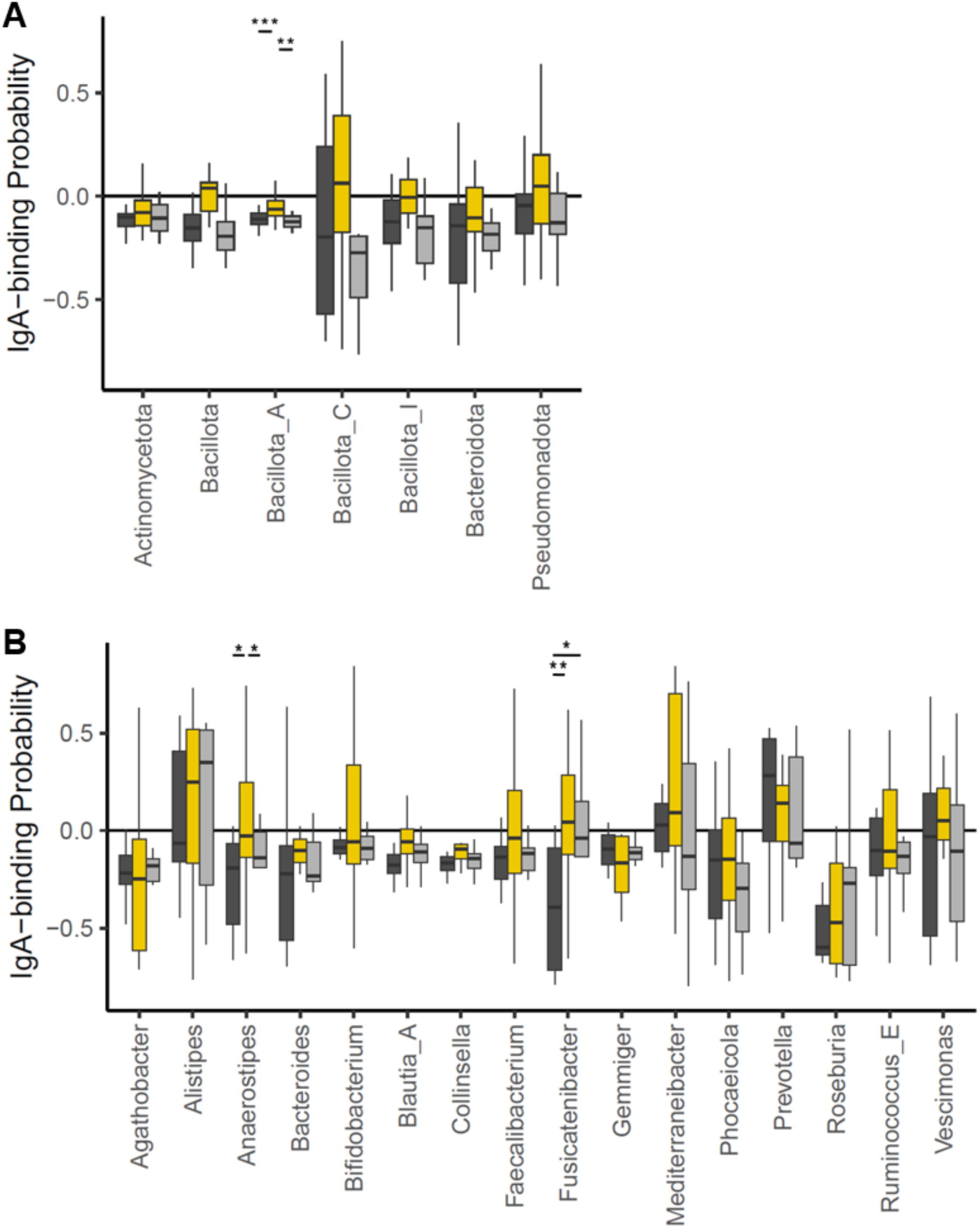
Comparison of SIgA binding Probability (SIgA-bP) of phyla (A) and genera (B) with a mean relative abundance >1%. Colors indicate allocation to respective cluster (dark grey = C1, gold = C2, light grey = healthy control (HC)). * - p<0.05, ** - p<0.01, *** - p<0.001, **** - p<0.0001.

## Notes

### Competing Interest Statement

The authors have declared no competing interest.

